# Fast processing of environmental DNA metabarcoding sequence data using convolutional neural networks

**DOI:** 10.1101/2021.05.22.445213

**Authors:** Benjamin Flück, Laëtitia Mathon, Stéphanie Manel, Alice Valentini, Tony Dejean, Camille Albouy, David Mouillot, Wilfried Thuiller, Jérôme Murienne, Sébastien Brosse, Loïc Pellissier

**Affiliations:** Department of Environmental System Science, ETH Zürich, 8092 Zürich, Switzerland; Swiss Federal Research Institute WSL, 8903 Birmensdorf, Switzerland; CEFE, Univ. Montpellier, CNRS, EPHE-PSL University, IRD, Montpellier, France; SPYGEN, Le Bourget-du-Lac, France; IFREMER, unité Écologie et Modèles pour l’Halieutique, rue de l’Ile d’Yeu, BP21105, 44311 Nantes cedex 3, France; MARBEC, Univ. Montpellier, CNRS, IRD, Ifremer, Montpellier, France; Institut Universitaire de France, IUF, Paris 75231, France; Univ. Grenoble Alpes, Univ. Savoie Mont Blanc, CNRS, LECA, Laboratoire d’Écologie Alpine F-38000 Grenoble, France; Laboratoire Evolution et Diversité Biologique (UMR5174), CNRS, IRD, Université Paul Sabatier, Toulouse, France

## Abstract

The intensification of anthropogenic pressures have increased consequences on biodiversity and ultimately on the functioning of ecosystems. To monitor and better understand biodiversity responses to environmental changes using standardized and reproducible methods, novel high-throughput DNA sequencing is becoming a major tool. Indeed, organisms shed DNA traces in their environment and this “environmental DNA” (eDNA) can be collected and sequenced using eDNA metabarcoding. The processing of large volumes of eDNA metabarcoding data remains challenging, especially its transformation to relevant taxonomic lists that can be interpreted by experts. Speed and accuracy are two major bottlenecks in this critical step. Here, we investigate whether convolutional neural networks (CNN) can optimize the processing of short eDNA sequences. We tested whether the speed and accuracy of a CNN are comparable to that of the frequently used OBITools bioinformatic pipeline. We applied the methodology on a massive eDNA dataset collected in Tropical South America (French Guiana), where freshwater fishes were targeted using a small region (60pb) of the 12S ribosomal RNA mitochondrial gene. We found that the taxonomic assignments from the CNN were comparable to those of OBITools, with high correlation levels and a similar match to the regional fish fauna. The CNN allowed the processing of raw fastq files at a rate of approximately 1 million sequences per minute which was 150 times faster than with OBITools. Once trained, the application of CNN to new eDNA metabarcoding data can be automated, which promises fast and easy deployment on the cloud for future eDNA analyses.

## 2 Introduction

Ecosystem governance and management require increasing the speed, accuracy and ease at which we can collect and process biodiversity data (Dornelas et al. 2019, Makiola et al. 2020), shifting the focus from expert monitoring towards high throughput data acquisition technology (Cordier et al. 2019). Conventional biodiversity monitoring approaches rely on the morphological identification of a limited number of taxa. Yet these visual surveys are labor intensive and require taxonomic expertise resulting in long delays between sampling and results (McGee et al. 2019). They miss many species that are either small, rare, cryptic or elusive (Iknayan et al. 2014) which leads to false negatives impacting ecological interpretations. Fortunately, our ability to rapidly generate inventories of whole species communities is growing with the emergence of environmental genomics, and specifically environmental DNA (eDNA, Bohmann et al. 2014, Thomsen and Willerslev 2015, Deiner et al. 2017, Cordier et al. 2020). All organisms living in an ecosystem shed tissue material, which can be detected by environmental DNA metabarcoding (Taberlet et al. 2012), offering an integrative view of ecosystem composition (Ficetola et al. 2008, Deiner et al. 2017). Coupled with high-throughput DNA sequencing methods, eDNA ‘metabarcoding’ can serve the rapid assessment and monitoring of biodiversity across all levels of life (from prokaryotes to eukaryotes) (e.g. Holman et al. 2021), with higher detection capacity and cost-effectiveness than traditional methods (e.g. Polanco Fernandez et al., 2021). The reads from high-throughput amplicon sequencing of eDNA are compared with reference barcode libraries allowing the establishment of taxonomic compositions directly from environment samples (Taberlet et al. 2012). Moreover, the resulting taxonomic lists can be linked with secondary information about taxa functional traits or phylogenetic position (Polanco Fernandez et al. 2021) and can be used to assess ecosystem functioning and health status (Cordier et al. 2020). While an increasing number of initiatives propose to use eDNA metabarcoding routinely and globally to monitor ecosystems (Berry et al. 2020), this would represent massive sequencing data which will require novel fast, accurate, and automated bioinformatic solutions.

As the laboratory molecular steps of eDNA metabarcoding have gained in efficiency (Shokralla et al. 2012, Thomsen and Willerslev 2015), the major bottleneck and technical challenge shifted from the development of efficient laboratory protocols to the processing of large metabarcoding data sets sequenced from eDNA into taxonomic lists (Dufresne et al. 2019). In particular, eDNA metabarcoding amplifies small DNA sequences (’barcodes’) typically of 80-300bp from the mitochondrial genome, from Illumina sequencing technology (Singer et al. 2019). This sequencing generates a huge quantity of small sequence reads that require fast and accurate bioinformatics processing to be interpreted (Ficetola et al. 2015, 2016). This bioinformatic processing includes several steps (the merging of the forward and reverse reads, demultiplexing, dereplicating, filtering by quality, removing errors) after which the clean sequences are assigned to a taxonomic label (Pagni et al. 2013, Dufresne et al. 2019, Marques et al. 2020a). Taxonomic assignment then transforms sequence reads from eDNA into lists of taxa that can be used by experts and policymakers for management decisions (Sepulveda et al. 2020) based on the detection of rare (Boussarie et al. 2018, Rojahn et al. 2021), endangered (Gold et al. 2020), or invasive species for example (Sepulveda et al. 2020). Yet, efficient algorithms transforming eDNA reads into accurate taxonomic lists are needed, potentially allowing parallel automatization on cloud infrastructure for a broad application of eDNA technology (Sato et al. 2018).

Compared with traditional bioinformatic approaches (Mathon et al. 2021), machine learning could increase the efficiency and capacity of eDNA reads treatment to assign taxonomic labels (Nugent et al. 2020). Machine learning has revolutionized object classifications in various biological applications from species identification on images (Grünig et al. 2021) to rare species distribution in habitats (Deneu et al. 2021). Taxonomic groups represent discrete classes that can be related to sequence features, including the composition and distribution of nucleobases within DNA sequences (Helaly et al. 2019, Busia et al. 2020). For example, k-mer summarises the counts of nucleotides within subsequences of length k and, in combination with machine classifications, have served for labelling sequences from bacteria, archaea, fungi and viruses (Piro et al. 2020). The association between k-mer features and taxonomic classification can be trained in a neural network from a reference genetic database (Piro et al. 2020), to predict the label of any new sequence. Alternatively, Convolutional Neural Network (CNN) can self-learn a broader range of spatially organised DNA base-motif features existing in the DNA sequences (Helaly et al. 2019). The neural structure subsets signal from restricted region of the input data known as the receptive field and can react to localized patterns in the sequence data. The numeric encoding of the four DNA bases allows the spatial placements of nucleotides to be interpreted by the CNN. In particular, Busia et al. (2020) developed a CNN, which trains a deep neural network to predict database-derived taxonomic labels directly from query sequences. Hence, preliminary use of machine learning with DNA sequence data shows their potential for taxonomic classification (Busia et al. 2020, Kopp et al. 2020), but this method has been mainly used so far to label longer amplicons (16S gene, up to 250bp, Busia et al. 2020). How it can be adapted for the taxonomic labelling of short sequence data from eDNA metabarcoding remains to be evaluated.

The most computationally costly step in the processing of eDNA metabarcoding is data cleaning (Mathon et al. 2021), and any computational gain from machine learning can only be reached if the CNN is robust to noisy sequencing data. Raw sequencing data can contain many errors including PCR substitutions or indel errors (Schirmer et al., 2015, Schnell et al., 2015, Taberlet et al., 2018). In existing eDNA bioinformatic pipelines the classification of DNA reads into taxonomic labels is applied after a long process of sequence processing and cleaning (Mathon et al. 2021) where only high quality reads are kept (Boyer et al. 2016). Thus data augmentation by artificially introducing variation in sequences within the reference database has been proposed to build a more robust CNN (Busia et al. 2020) allowing the processing of raw Illumina sequencing data. For example, Busia et al. (2020) applied a data augmentation in the training phase by adding between 0.5 and 16% of mutations by switching DNA bases randomly. While it was shown that moderate artificial noise renders the network more robust to potential sequencing errors, setting an excessive value decreases the CNN performance (Busia et al. 2020). In addition, a robust CNN could be trained to tolerate the PCR tags and the remaining primers as present in raw metabarcoding data, but these aspects remain largely unexplored. A robust CNN could then be used to process and identify the sequences found within an entire metabarcoding (fastq) file independently of assigning the identified reads to each sample. If reliable, such a pipeline would be a major addition to existing tools to process the exponentially growing quantity of available metabarcoding data in eDNA analyses.

Here we evaluated the training of a CNN on a reference database of genetic sequences and its ability to rapidly and accurately process raw eDNA metabarcoding data. More precisely, we aim to adapt CNNs to the short sequences produced by eDNA metabarcoding and test whether accuracy and speed of CNNs are comparable to that of OBITools, a widely used bioinformatic pipeline (Boyer et al. 2016). As a case study, we used one of the largest standardized eDNA data set currently available for fishes corresponding to a multi-year campaign effort to sample the tropical South American rivers of French Guiana (Murienne et al. 2019). This eDNA data set is associated with a quasi-exhaustive reference database covering most of the known species of the region for the “teleo” region of the 12S rRNA mitochondrial gene (Cilleros et al. 2019, Coutant et al. 2020). The freshwater ecosystems of French Guiana are among the most species-diverse ecosystems for riverine fishes globally (Albert & Reis 2011), and among the least human impacted rivers on earth (Su et al. 2021). A demonstration of a good performance in such diverse ecosystems would provide a robust test for application in other simpler ecosystems globally. Within this general processing framework and using this case study, we ask the following questions: (i) How does a CNN approach perform in the training of eDNA sequences classification for labels of the reference database; (ii) How robust is the classification of a CNN applied directly to the raw Illumina metabarcoding short sequences; (iii) How do a classical metabarcoding pipeline and our CNN approach compare with the pre-existing information about biodiversity composition within two river catchments with a long history of traditional sampling effort? Accurate performance of the CNN across all these steps would allow benchmarking the application of machine learning for the processing of Illumina sequence data and would open a major bottleneck in biodiversity analyses.

## 3 Material and Methods

### 3.1 eDNA data collection and reference database

As a test data set we used data collected in French Guiana, a *c*. 80,000 square kilometers South American territory almost entirely covered by dense primary forest (Supplementary Fig. 6). Equatorial climate associated with abundant rainfall created a dense hydrographic network made of six major watersheds and several coastal rivers that host a highly diverse fish fauna with at least 368 strictly freshwater fish species (Le Bail et al. 2012). eDNA field collections have been initiated in 2014 and have continued until 2020. We sampled over 200 sites (see Murienne et al. 2019 for detail), where we filtered 30 liters of river water across a flow filtration capsule using a peristaltic pump. For the purpose of this study, we analyzed only the filters collected in the Maroni and Oyapock rivers.

In each site we collected from one to ten filtration capsules but in most sites two capsules were used (2×34 litres) using the protocol described in Cantera et al. (2019) and Coutant et al. (2020). A peristaltic pump (Vampire sampler, Burlke, Germany) and disposable sterile tubing were used to pump the water through the encapsulated filtering cartridges (VigiDNA 0.45 μM, SPYGEN). The input part of the tube was held a few centimeters below the surface in rapid hydromorphologic units to allow a better homogenisation of DNA in the water column. When filters began to clog, the pump speed was decreased to avoid material damage. To minimize DNA contamination, the operators remained downstream from the filtration either on the boat or on emerging rocks. After filtration, the capsules were filled with a preservation buffer and stored in the dark at room temperature for less than a month before DNA extraction. The 12S rRNA “teleo” gene fragment (Valentini et al. 2016) was amplified by PCR and sequenced on an Illumina platform generating an average of 500,000 paired-end sequence reads per sample. The DNA extraction, amplification and sequencing protocol are described in Cantera et al. 2020.

We generated an eDNA reference database by combining fish specimens caught using various fishing gears. These data were complemented by fish collections carried out by environmental management agencies (DGTM Guyane, Office de l’eau Guyane, Hydreco laboratory), fish hobbyists (Guyane Wild Fish), and Museum tissue collections (MHN Geneva). Although rare for Guianese fishes, existing sequence data from online databases (Genbank, Mitofish) were also included. We extracted and sequenced the 12S ribosomal gene on the species collected. Our local reference database has improved over the years (Cilleros et al. 2019, Cantera et al. 2019) and now covers over 368 species out of 380 estimated in the region, so an almost full coverage which remains exceptional regarding the main gaps globally (Marques et al. 2020b).

### 3.2 OBITools bioinformatic pipeline

As a standard processing pipeline we selected OBITools (http://metabarcoding.org/obitools, Boyer et al., 2016) which is commonly used in many eDNA metabarcoding studies (Bylemans et al. 2018, Li et al. 2021, West et al. 2021). Reads from the sequencing were processed following Valentini et al. (2016). In short, the forward and reverse reads were assembled using `illuminapairedend` with a minimum score of 40 and retrieving only joined sequences. The reads were then assigned to each sample using `ngsfilter`. A separate data set was then created for each eDNA sample by splitting the original data set into several files using `obisplit`. After this step, each sample was analyzed individually before generating the taxonomic list. Strictly identical sequences were clustered together using obiuniq. Sequences shorter than 20 bp were excluded using `obigrep`. Then we ran `obiclean` within each PCR product for clustering. We discarded all sequences labelled as ‘internal’ corresponding most likely to PCR substitutions and indel errors. Taxonomic assignment of the remaining sequences was performed using `ecotag` with the custom genetic reference database relevant for the eDNA samples. Finally we applied an empirical threshold to account for tag-jumps and spurious errors.

### 3.3 Reference data augmentation and training data set

The reference database has a full species coverage but the number of DNA replicate sequences for each species is limited as there are only 683 sequences for 368 species. This makes training a CNN challenging for several reasons. The number of sequences per species is not balanced, there are not enough sequences to capture the entire inter- and intraspecific variation, and the noise from the sequencing process is not accounted for. To balance the data set using data augmentation procedures we over-sampled the underrepresented species before training. To increase the sequence variation, we implemented an inline sequence mutation step similar to Busia et al. (2020). During each training epoch all sequences were randomly mutated. We added between zero and two random insertions and deletions each as well as noise in the form of a 5% mutation rate. This procedure further reduces overfitting as no training sample is likely to be repeated twice. For the evaluations, we either added no augmentation or 2% noise and singular insertions and deletions as we expected the PCR amplification and sequencing to be better than the 5% noise considered during the training.

For the direct application on the raw reads, another data transformation was required. All sequences processed in an Illumina machine retain the selected primers and were tagged with 8bp long tags. During the sequencing two bases from the plate attachment sequence were often read as well. We therefore pre- and appended the forward and reverse primers, and the combined tags and attachment bps to the sequences from the reference database. Specifically we added 10 bp of unknown bases to each reference sequence, represented by the IUPAC ‘N’ code. This shifted the sequences to a position in the training input similar to where they would occur in the Illumina data. While there is a canonical read direction for DNA the read directions during the sequencing randomly occurs in either directions. Therefore we added the reverse complement of all sequences to the final data set. As the last step we truncated all sequences to 150bp for the read length as fixed by the field metabarcoding data.

### 3.4 CNN training and evaluation using split sampling

We investigate the performance of a Convolutional Neural Network (CNN) approach trained on the reference database at the species level. To encode DNA sequence information, each canonical base (A, C, T, G) and IUPAC ambiguity codes are translated to an appropriate four-dimensional probability distribution over the four canonical bases (A, T, C, G) including uncertain base reads (e.g. W and S). For example ‘A’ becomes [1,0,0,0] or ‘W’ equals [0.5, 0, 0, 0.5]. The neural network was designed and optimized through a series of tests that allow for the optimal set of correct DNA features to be selected. In particular, we explored an exhaustive number of model sizes, including one to three layers of 2d (depth-wise) separable convolutions with 4-16 filters each, one to three fully-connected layers with varying numbers of neurons each, and a softmax activated output layer which produces a probability distribution over all possible taxonomic labels. We applied dropout regularization and used leaky rectified-linear activation for all but the last layers.

We used TensorFlow (Abadi et al., 2015) to train the CNNs with all the aforementioned data augmentations. Due to the sparse dataset, we characterized and evaluated the performance of the neural networks by using three different methods. First, we applied a cross-validation with random split-sampling from the reference database. This establishes a proper separation between the training and validation data, but less than half the species in the reference dataset have two or more sequences resulting in a reduced range of species to be included. Here only 156 out of 368 possess more than two unique sequences are were considered for the split dataset. Next, we trained several networks on the full reference data set with all 368 species and validated them on the original non-augmented reference data. Finally, we derived more synthetic data from the reference sequences similar to the training augmentations and evaluated them with the chosen network. We evaluated whether there are systematic errors in the CNN performance. We further investigated whether a binarisation threshold, where we require a probability of the most likely prediction to be above a certain value, can improve the classification performance. As we privilege the absence of errors, i.e. less false positives, above the presence of correct predictions we evaluated the effects of such a binarisation threshold using the F-beta measure that allows for a weighted trade-off between these errors. As such we chose a small beta of 0.3 to heavily discourage false positives at the cost of discarding more correct results.

### 3.5 CNN application on demultiplexed and cleaned samples

We tested the best trained CNN on the curated eDNA reads after the application of the main cleaning steps of the OBITools pipeline. In particular, from the Illumina raw output, the forward and reverse reads were assembled using the `illuminapairedend` algorithm from the OBITools package, after which only high quality reads were kept, and demultiplexed across the different eDNA samples. We applied the best trained CNN at the species level on these curated eDNA samples. We compared the taxonomic assignments performed by the CNN to classic assignments using ecotag from OBITools. We evaluated and applied different thresholds for accepting species detection as a way to remove spurious errors and wrong assignments (0, 5, 10, 25, 50, 75 and 100 reads in at least one PCR replicate). For each eDNA PCR replicate, filter, and for the whole rivers we ranked the taxonomic groups by the number of reads recovered by each method and performed a Kendall rank correlation. We ran one rank correlation per eDNA sample and reported the median rank correlation across all samples. In addition, we compared the presence-absence using the kappa statistic measuring general agreement of the methods for each sample. We reported the median percentage and median kappa values across samples. Then, across all eDNA samples, we correlated the species richness obtained via CNN with that obtained with OBITools. Each analysis was performed at three different scales: the PCR replicate, the filtration capsule and the river. Finally, we ordinated species composition of each filtration capsule for both methods using a Principal Coordinate Analysis (PCoA) to compare differences in recovered compositions among methods.

### 3.6 CNN application on the raw illumina sequences

We applied the best CNN directly on the raw outputs from the Illumina sequencing, where we omitted all the preprocessing steps from OBITools. The CNN was expected to learn how to ignore the primer as it is constant for all presented sequences. Furthermore, the output sequences from the Illumina sequencer were fixed in length (150bp) so we fixed the input width of the CNN to this size. We systematically zero-padded or truncated the input sequences to this length during training, evaluation, and application. After the training with the reference database and the application on fastq, we developed a custom code for the fast demultiplexing of the reads. By focusing on the tag information in the first few positions of the sequence and not considering read errors in tags, we reduced the demultiplexing to a few simple look-ups in a hash table (currently 5), therefore reducing computational time with limited information loss. As in the previous test, we obtained a list of taxonomic assignment for each eDNA sample, which can be compared to species composition obtained with the OBITools pipeline. We further applied a threshold approach obtaining predicted composition per sample for any threshold tested. As done previously, for each eDNA sample, we ranked the taxonomic groups by the number of reads recovered by each method and performed a rank correlation. We reported the median rank correlations across all the eDNA samples. In addition, we compared the presence-absence at the species level using the overall kappa statistic measuring general agreement of the methods for each sample. We further evaluated whether differences between methods were more frequent in specific taxonomic families than others. Then, across all eDNA samples, we correlated the species richness obtained via CNN compared to that obtained with OBITools, and ordinated species composition of each filter for both methods on a PCoA. We evaluated the change in accuracy between the CNN applied to curated reads compared to raw fastq files.

### 3.7 Validation with existing biodiversity knowledge on the region

We finally compared the species composition recovered in the eDNA samples by CNN and OBITools to the species, genus and family check lists of each river catchment. Species lists for each catchment were obtained based on an updated version of the catchment scale species lists provided in Le Bail et al. (2012). From this list, we updated the taxonomy and added novel occurrences of known species based on fish catches by several research and management organizations (see above). Only collected specimens with validated taxonomy were considered to update this list, and detections using eDNA were not considered. We specifically quantified the number of matching species, false presences and false absences from each method, taking the checklists as references.

## 4 Results

### 4.1 CNN training and evaluation with split sampling

The exploration of CNN complexity showed that larger networks did not necessarily produce better results, indicating low overfitting and that a CNN of moderate complexity could learn the full structure contained in the reference data (Supplementary data set 1). The training and evaluation of the CNN with split sampling only considered 156 species (out of 368) which had more than two unique sequences. The optimal CNN consisted of one convolutional layer of 4 filters with a 7×4 extend, 3 dense layers with 128 neurons each, and the 156 neurons wide output layer. On the training data, the networks achieved 92% accuracy with little differences between the networks trained on the clean and the augmented (”raw”) sequences. When applying the CNN to held out data (316 sequences from the 156 species), we found a 91% accuracy on the clean data and 89% on the augmented data. Using an optimized 0.9 binarisation threshold with the F-beta metric, the accuracy rose to 98% at the cost of discarding 16% to 26% of the data respectively. We then used the entire data set in the training, including the 368 species, and repeated the analyses for clean and augmented data. The optimal model size was similar to the previously chosen networks, with a single convolutional layer of 4 filters with a 4×7 extend, followed by 2 dense layers each 384 neurons wide. With these networks, training accuracy was similar to that of internal evaluation, with 92%. Validating the networks on the clean and augmented reference data yielded higher accuracy at 96% and 94% respectively. With a binarisation threshold of 0.9, the accuracy rose to 99% for both the clean and augmented networks at the cost of rejecting 9% to 13% of all sequences evaluated (Fig. 1). We use a binarisation value of 0.9 for all further evaluations.

**Figure 1:**
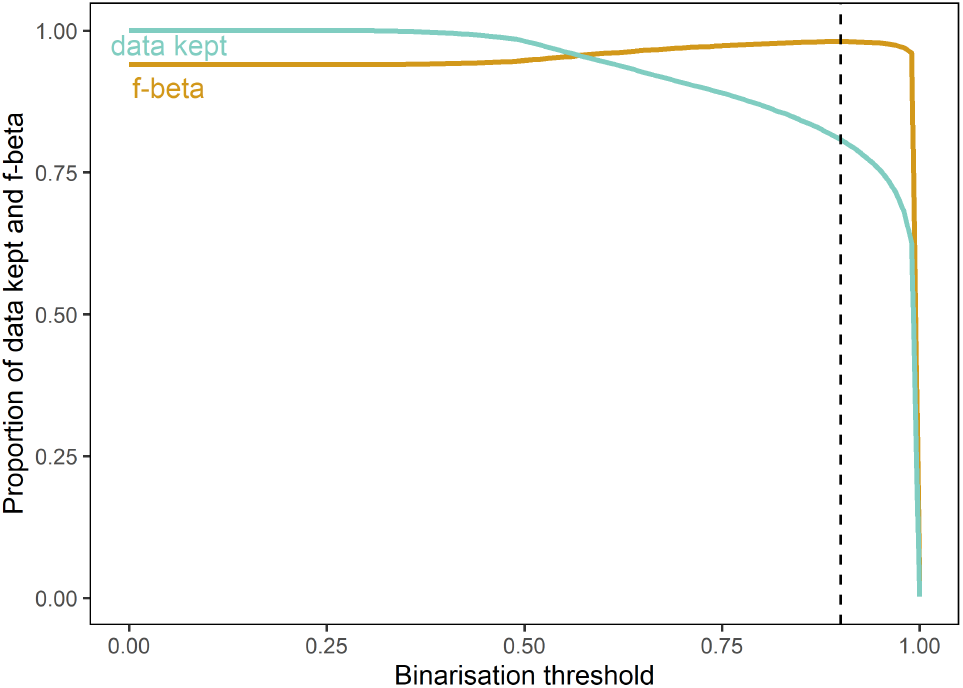
F-beta measure (orange) based on the predictions of the CNN on synthetic data after the training phase, for each binarization threshold value. Proportion of data discarded (blue) for each binarization threshold. The dashed vertical line indicates the threshold of 0.9, providing the highest F-beta value with minimum data discard.

### 4.2 CNN application on the raw and cleaned eDNA data set

The Kendall Tau-b correlation between the number of reads in OBITools and the CNNs increased when a more stringent threshold on the minimum number of reads per species in one PCR replicate was applied and stabilised with a threshold of 50 per species and minimal threshold were considered (Fig. 2). We applied a threshold of 50 reads per species within a PCR replicate in the following analyses. At the PCR replicate level, the correlation between the CNN applied on raw reads and OBITools was significantly higher than the correlation between the CNN applied on clean reads and OBITools, but that relation was inverted at the filter level. When applying the CNN on raw reads we found a median kappa value of 0.93 across all eDNA filters (range 0.79-0.99), with a slight difference toward more species predicted by the CNN (median species number 63), than OBITools (median species number 56) (Fig. 3). We found a median Kendall Tau-b rank of 0.77 across all the eDNA filters (range of Kendall Tau-b values 0.22-0.94). When applying the CNN on the clean reads after OBITools, we found a significantly higher median correlation (Kendall Tau-b 0.84, range=0.2-1), and median kappa value (0.96, range 0.83-1.0). The PCoA also indicated a better match of filter taxonomic composition between the CNN applied on clean data and the OBITools (Fig. 4) (A) than from the CNN applied on raw data and OBITools (B).

**Figure 2:**
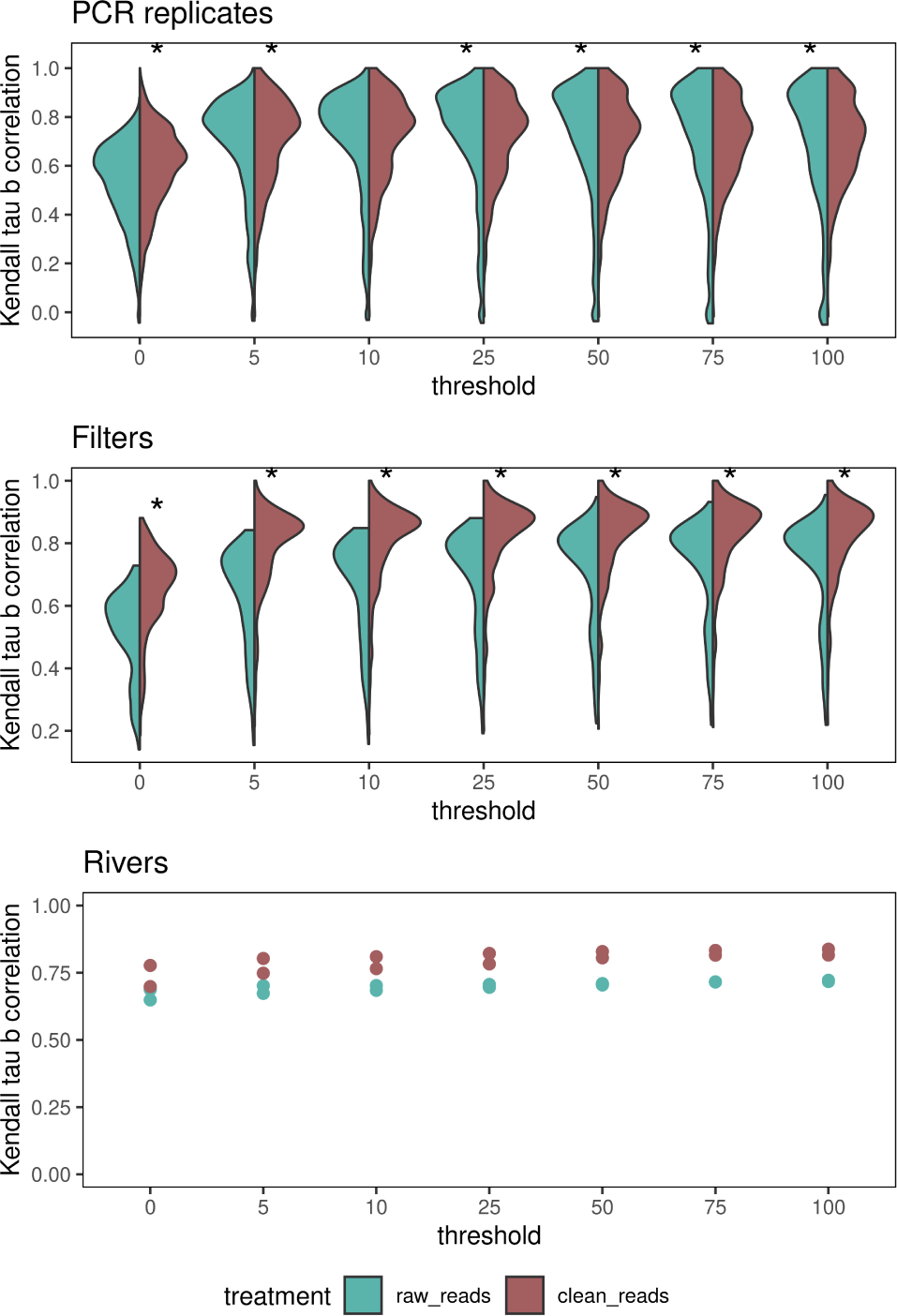
Kendall Tau-b correlation coefficient between the outputs of the CNN and OBITools. Left side of the violin plots (blue) represent correlation values between OBITools and the CNN applied on raw reads. Right side of the violin plots (red) represent correlation values between OBITools and the CNN applied on clean reads. The x-axis represents the threshold of minimum read number per species to be considered present. Stars represent a significant difference between the correlation from the CNN applied on raw reads or clean reads. The analysis was made at three levels: PCR replicates (top), filters (middle) and rivers (bottom).

**Figure 3:**
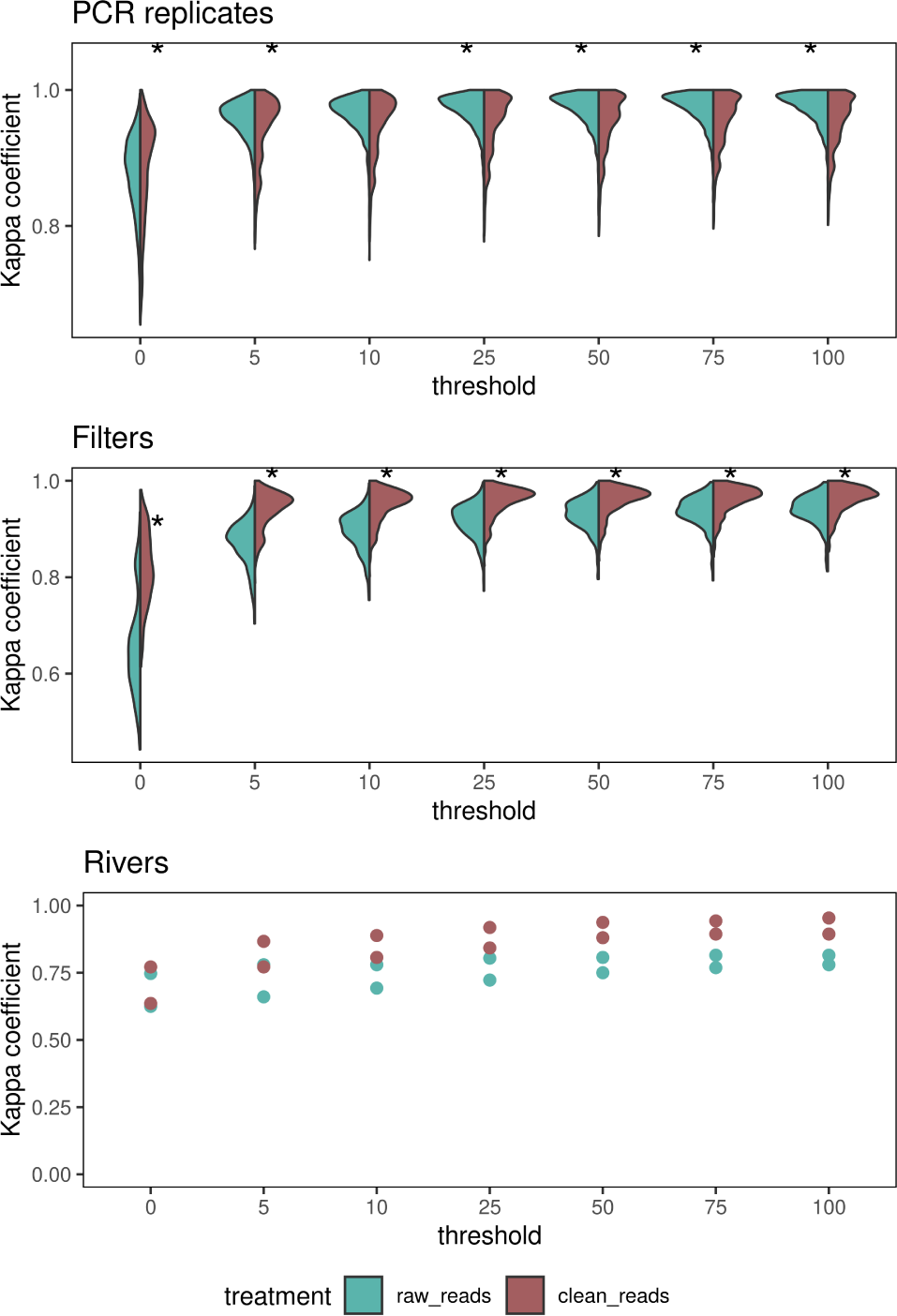
Kappa correlation coefficient between the outputs of the CNN and OBITools. Left side of the violin plots (blue) represent correlation values between OBITools and the CNN applied on raw reads. Right side of the violin plots (red) represent correlation values between OBITools and the CNN applied on clean reads. The x-axis represents the threshold of minimum read number per species to be considered present. Stars represent a significant difference between the correlation from the CNN applied on raw reads or clean reads.

**Figure 4:**
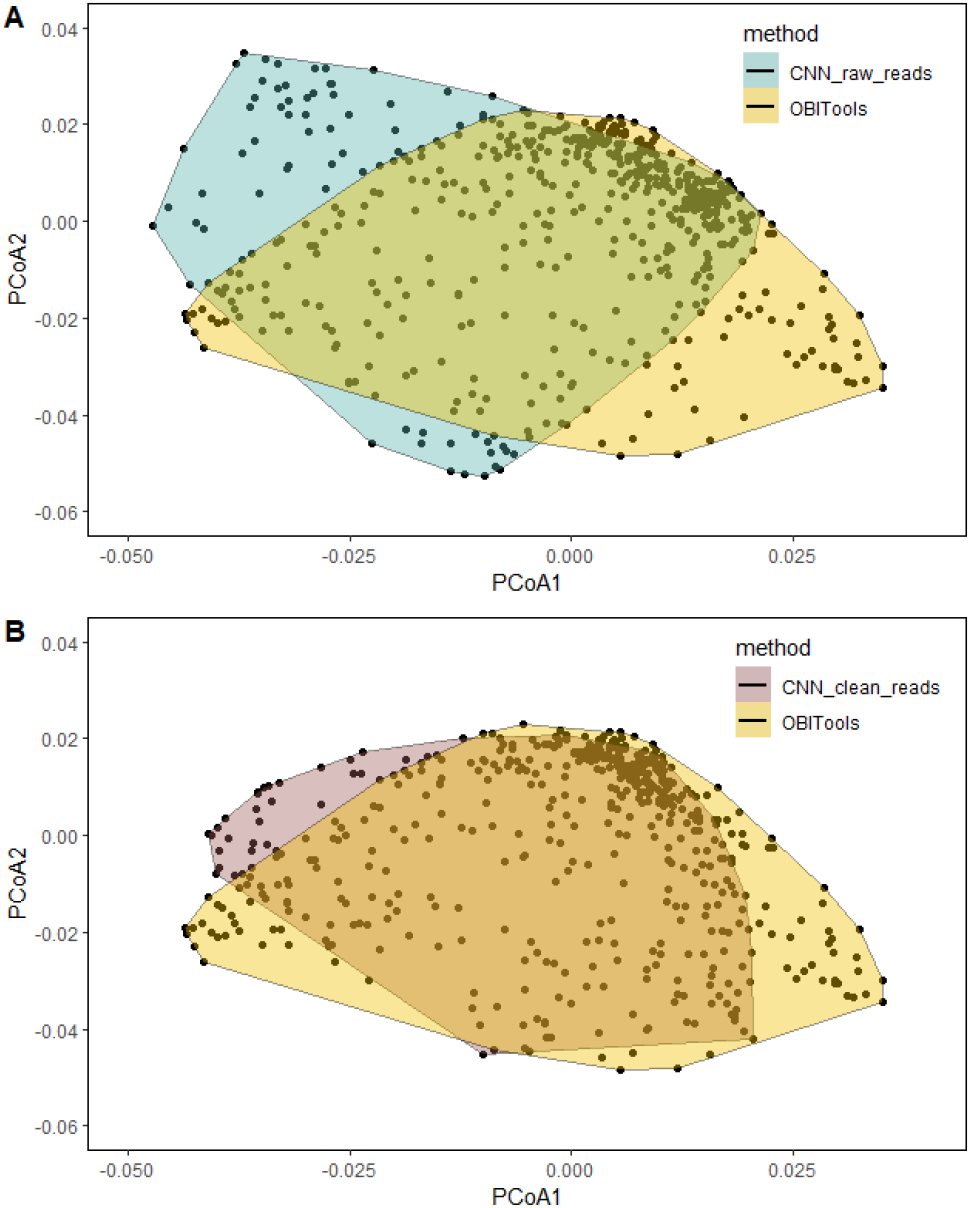
Principal Coordinate Analysis (PCoA) on species composition similarity between filters. A, ordination of filter species composition similarity in the outputs of the CNN applied on raw reads (blue) and in the outputs of OBITools (yellow). B, ordination of filter species composition similarity in the outputs of the CNN applied on clean reads (red) and in the outputs of OBITools (yellow). Similarity matrices were built with Bray-Curtis distances on reads abundance per species per filter.

### 4.3 Validation with known biodiversity in the region

The data synthesis across historical fish surveys yielded 351 species in the Maroni and Oyapock combined, among which 293 are present in the reference database and thus potentially detectable with eDNA. We used a threshold of 50 reads for a species to be considered as present in both pipelines. For both rivers combined, the CNN applied on raw reads assigned 319 species, among which 264 are known from historical records, while 55 were never recorded (Fig. 5a). The CNN and OBITools detected 274 species in common, while the CNN retrieved 21 species known from historical surveys in these rivers that were not retrieved with OBITools, but identified 24 species not known from the synthesis nor identified by OBITools. The species detected only by the CNN belong mainly to the Loricariidae, Cichlidae, Characidae and Callichthyidae families. The 23 species known from historical records and not detected by either eDNA method belong mainly to the Loricariidae, Characidae, Apteronotidae and Anostomidae families. The two species detected only with OBItools are from Cichlidae and Aspredinidae families (Fig. 5b). The CNN applied on clean reads detected 293 species, of which 254 are present in the Maroni and Oyapock synthesis, 276 were common with the outputs of OBITools, 9 were found only with CNN and synthesis, and 8 were found only by the CNN.

**Figure 5:**
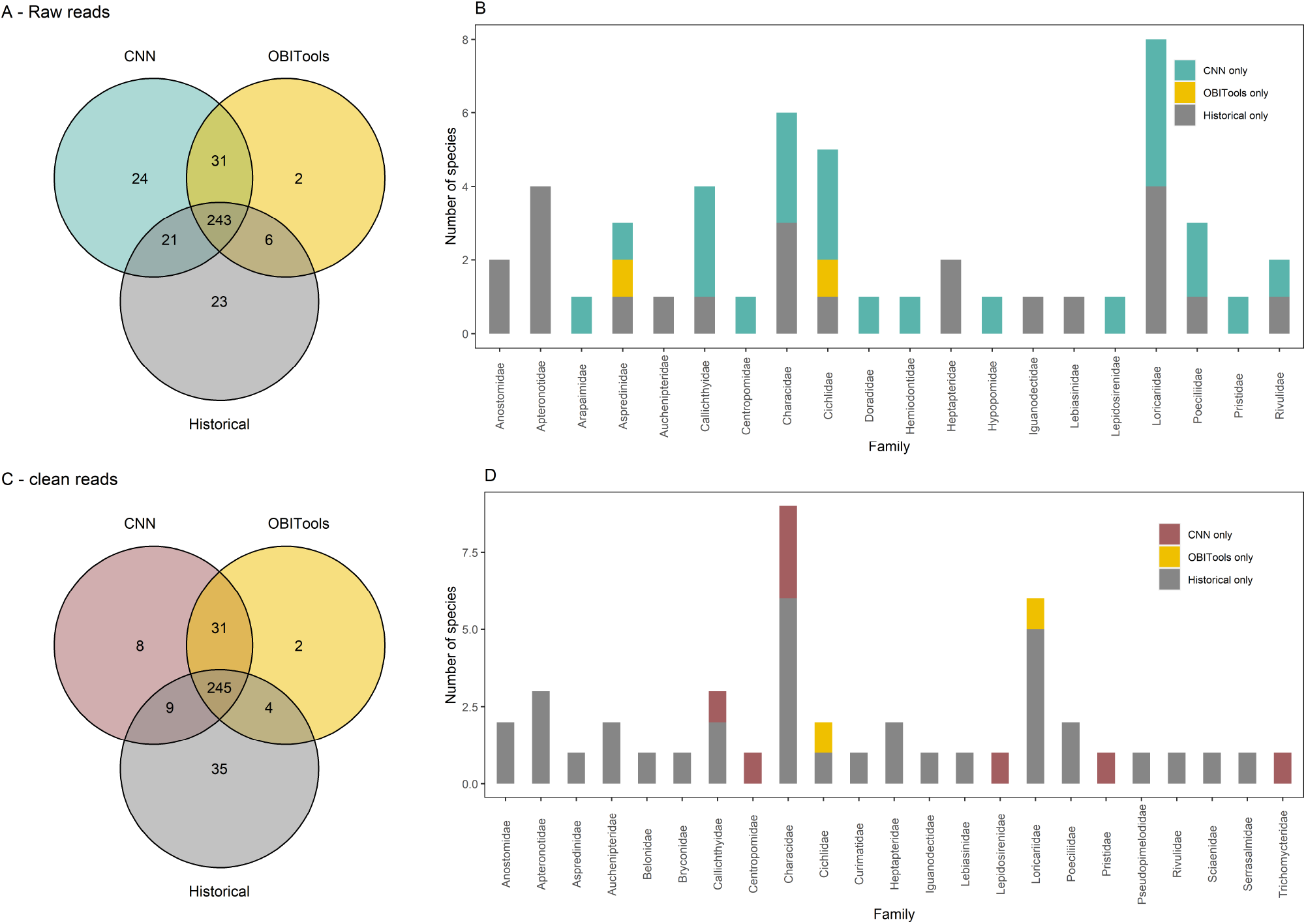
Species detections with the CNN, OBITools and in historical records, in the Maroni and Oyapock rivers combined. A, overlap of species detections between the CNN applied on raw reads (blue), OBITools (yellow) and historical records (grey). B, number of species per family, detected with only one method (CNN on raw reads, OBITools or historical records). C, overlap of species detections between the CNN applied on clean reads (red), OBITools (yellow) and historical records (grey). D, number of species per family, detected with only one method (CNN on clean reads, OBITools or historical records)

In the case of OBITools, 282 species were detected, among which 249 are known from historical synthesis, while 33 were never recorded in the Maroni or Oyapock (Fig. 5c). The species detected only by the CNN belong mainly to the Characidae family. The species known from historical records not detected by either eDNA method belong to the Loricariidae, Characidae and Apteronotidae families. The two species detected only with OBItools are from Loricariidae and Cichlidae families (Fig. 5d). The same analysis at the scale of each river provided similar results(Supplementary Fig. 7-8). Hence, while both methods detected species not found in historical sampling, the CNN is recovering generally more species than the OBITools, which could correspond to both new true observations or commission errors. The CNN applied on raw reads retrieved a higher number of species that were not recorded from historical records nor found with OBITools. For the Maroni, the CNN applied on raw reads and the CNN applied on clean reads retrieved 232 species in common, while 48 were found only with the raw reads and 16 only with the clean reads. For the Oyapock, 185, 66 and 18 species were found in common only with the raw reads and only with the clean reads, respectively (Supplementary Fig. 9).

### 4.4 Computation time

For the CNN, we have to distinguish two computational efforts, which can be measured independently. First, training the networks needs to be performed once per reference database, and second, the application on field data. Training a network on the augmented and complete reference database currently takes around 10 minutes on an Nvidia Titan RTX GPU. Training a network on the clean reference database is faster and takes 6 minutes on the same GPU. The training and application time is dependent on the size of the input data and network size. Applying the network to field data processes around 1 million input sequences per minute compared to only 20’000 input sequences per minute for OBITools. A large part of the computational time for the OBITools pipeline is dedicated to the alignment (up to 80%) and demultiplexing (up to 15%) steps. By training and applying a convolutional neural network directly on raw reads, we can sidestep this issue completely and achieve significantly faster processing times and lower power consumption at the expected cost of a slightly lower accuracy in the results overall.

## 5 Discussion

An emerging technical issue with eDNA application for biodiversity monitoring is that, as sequencing technology becomes cheaper and more affordable, the quantity of data to be processed induces a major computing bottleneck. Our study demonstrates the application of a Convolutional Neural Network (CNN) to process short eDNA sequence reads directly from raw sequencing Illumina outputs. In only a few minutes, the software can transform a raw fastq sequence data set in a species list associated with each eDNA sample collected in the field. The processing time of the CNN contrasts with standard bioinformatic pipelines, which can require significant computing time to process the raw reads into taxonomic lists (Mathon et al. 2021). We further show that the CNN approach delivers species composition roughly comparable to OBITools, and historical knowledge. While limitations remain in the first application presented here, future developments are likely to improve the speed and accuracy at which CNNs can translate raw metabarcoding data into taxonomic lists. As the coverage of the reference databases may improve in a near future (Marques et al. 2020b), CNNs could offer an efficient way to revisit stored metabarcoding data and increase biodiversity knowledge on previously sampled sites. Together, machine learning offers new possibilities for the taxonomic assignment of short DNA sequences and transform fast collected eDNA data into interpretable taxonomic-based indicators for the use of stakeholders (Cordier et al. 2020, DiBattista et al. 2020, Sepulveda et al. 2020).

The CNN offers fast and accurate processing of a large number of sequences applied directly on the raw reads from the Illumina outputs. In classical bioinformatic pipelines, the processing from raw sequence reads to taxonomic identifications include seven steps (paired-end reads merging, demultiplexing, dereplication, quality filtering, removal and correction of PCR/sequencing errors, and taxonomic assignment) expected to be essential to generate high quality results from metabarcoding studies, but which can be computationally demanding (Bonder et al. 2012, Calderón-Sanou et al. 2020) and challenging to articulate (Marquez et al. 2020a). We show that the CNN can embed all these steps in a single process applied directly on the raw Illumina reads when the CNN is trained to handle noisy data. Moreover, for relatively short eDNA markers (e.g. 60bp for the ‘teleo’ marker as used here), merging paired-end reads is not necessary, which leads to a significant computational gain (Mathon et al. 2021). While offering results roughly comparable to those of OBITools, the CNN decreases the processing time of the whole dataset analysis by a factor of around 150. In a recent comparison, Barque (https://github.com/enormandeau/barque) combined with a fast demultiplexing module, allowed to proceed over 15Mio reads in 30 minutes, while it took 17 hours for OBITools V1 (Mathon et al. 2021). Assuming the same rate of our CNN of 1 Mio read per minute on this data set the application of the CNN would remain two times faster than the fastest existing bioinformatic pipeline in a single model (Mathon et al. 2021). Our study presents a first successful adaption of CNN to the processing of eDNA metabarcoding data, but we foresee several avenues of optimization to gain speed and accuracy, making it a promising tool for scaling-up biodiversity inventories via eDNA (Berry et al. 2019, Ruppert et al. 2019).

The training of CNN allows an efficient adjustment to the reference database, avoiding the need to explore a large number of parameters and arbitrary thresholds as in classical bioinformatic pipelines. Existing bioinformatic pipelines contain a variety of modules (i.e. QIIME2, DADA2, Vsearch), each with its own set of parameters (Bolyen et al. 2019, Callahan et al. 2016, Rognes et al. 2016). Selecting the appropriate modules and parameters requires advanced knowledge of the functioning of the program since changes in those parameters can considerably modify the outputs (Flynn et al. 2015, Bonder et al. 2012, Brown et al. 2015). The absence of an appropriate and automated method for parameter optimization (Alberdi et al. 2018) often challenges the use of those pipelines by non-specialists. In contrast, the application of a CNN only includes a first step of training, where the optimisation of the network is near automated, and two independent steps for applying the CNN and de-multiplexing the reads to reach to final taxonomic outputs per sample. During the learning step of a CNN, only three parameters have to be set by the user: the network size (number of layers, filters and units), the learning rate and the augmentation values. During the application step, only two parameters have to be set by the user, the binarization threshold and the minimum number of reads per sample to be considered. We expect that those steps can be near automated within a user-friendly software as developed in other machine learning application (Thuiller et al. 2009). Given the relative ease in the training and application of CNN, it could be transformed into an application with a user-friendly interface and only a minimum of actions required from the user. Hence, CNNs could make eDNA metabarcoding data processing accessible even to less trained users. And they could, for example, be combined with the minion sequencer to provide an instantaneous view of the biodiversity.

A CNN trained on a complete reference database produced species composition congruent with the outputs of a popular bioinformatic pipeline, but showed a tendency to predict more species than those of OBITools and historical records. Compositional differences in the outputs of pipelines have already been highlighted (e.g. Brandt et al. 2021) and mainly resulted from the detection of several false positives and false negatives (Mathon et al. 2021). With a binarization threshold of 0.9 optimized during the training, we found congruent but slightly diverging results between OBITools and the CNN when applied to both the raw and clean reads. While the CNN and OBITools shared most of their recovered species, each method detected a few species not detected by the other (Fig. 5). However, the CNN showed a general tendency for overprediction compared to OBITools and historical records, especially when applying it directly on the raw sequencing data. Using the historical records as a baseline, the CNN on clean reads reduces the detection of species only found with the CNN, without decreasing the number of species shared with OBITools or historical records, suggesting false positives result from noisy inputs. Specifically, the CNN on raw reads detected a higher number of species from Loricariidae, Cichlidae and Characidae families that are not found with OBITools, which can come from sequencing errors that are not denoised by the CNN. In the case of the Cichlidae family, the short barcode we use is known to be poorly resolutive (Taberlet et al. 2018), with a lot of species sharing the same sequence (Polanco et al. submitted), and our CNN cannot perform well in this situation like all other pipelines. Moreover, Loricaridae and Characidae are the two most speciose families from the Guianese fish fauna, with more than 50 species per family (Le Bail et al. 2012), and with several new species occurrences recorded each year in Guianese rivers (e.g. Brosse et al. 2019). Those two families, together with Cichlidae are also known to host cryptic and still unnamed species, as shown by Papa et al. (2020) for the Maroni. This could also contribute to the unattended species detection. Finally, we found that the correlation between OBITools and CNN was lower at the level of the filter, than at the level of the PCR replicates when applying the CNN on raw reads, but higher at the filter level when applying the CNN on clean reads. Hence, appropriately combining the PCR replicates could confer more robustness to the final outputs of the CNN. Refinement of the network could be added, so that the detection across multiple PCR replicates can be used to compute the final likelihood.

Our study proposes a first application of CNN to eDNA metabarcoding data, but several improvements are required before broad scale future applications to eDNA big data. The current CNN is learning from species class and is forced to assign the sequences to that taxonomic level. Thus, when presented with conflicting sequences, the network might assign all of them to a single species, or may split the probabilities across several species, which could be then discarded given the use of the 0.9 binarisation threshold. In contrast, in case of conflict, OBITools can assign sequences to higher taxonomic levels, allowing to keep information related to these species with identical sequences. In this case study, we were in the ideal situation where the reference database is almost complete for the territory and the CNN can be improved to be able to handle incomplete reference database and be able to assign the read to other taxonomic levels or to an unknown class, rather that forcing a species-level identification and relying on the binarisation threshold to reject unknown sequences. We could also improve the CNN by implementing more stringent filters that would reduce the number of false detections and prediction errors. For instance, some filters for tag jump handling included in previous pipelines for eDNA matabracoding for fish (e.g. Cilleros et al. 2019) are not included here. The CNN we present in this study is a first proof of concept for the application of machine learning on eDNA metabarcoding data, but further improvements are possible, especially in regards to dealing with inconsistent reference databases, and implementation as a user-friendly interface.

### 5.1 Conclusion and perspectives

We demonstrate that we can use machine learning to increase the speed and decrease the energy consumption for processing eDNA metabarcoding data with good accuracy on clean reads and slightly lower accuracy on raw reads. The highest computation time for the CNN is on the training phase, but once trained, the CNN can be used as a computationally efficient tool for the application in the cloud facilitating the analyses of the mass of eDNA data collected in future biodiversity surveys. eDNA data are now produced at an exponentially increasing rate. By its easy application due to the reduced number of processing steps and the automated learning of best suited parameters, a CNN approach contrasts with other widely-used bioinformatic pipelines. Our work paves the way towards computationally efficient and user-friendly online processing pipelines that will allow the democratisation of bioinformatic analyses of eDNA samples. Our work is a major complement to the recent development and standardisation of eDNA in the laboratory, which together allow for extending the use of eDNA in community ecology and biogeography even for poorly known ecosystems or lineages (Juhel et al. 2020), and install eDNA as a standard monitoring tool (Jarman et al. 2018). It also reinforces its initial goal of quick and efficient application. We expect that the results from this study will be scaled up to become a major toolkit for ecological analyses of eDNA data possibly associated with a cloud infrastructure and parallel computation on GPU.

## Acknowledgments

This work benefited from Investissement d’Avenir grants managed by the French Agence Nationale de la Recherche (CEBA: ANR-10-LABX-25-01; TULIP: ANR-10-LABX-0041) and ANR grant (DEBIT: ANR-17-CE02-0007-01).

## Author contributions

B.F. and L.P. conceived the idea, study design, and analytic methods. B.F. developed the neural networks and ran the computational study. J.M. and S.B. collected the eDNA samples. A.V. and T.D. did the eDNA laboratory and data preparation work. B.F. and L.M. analysed the results. L.P., L.M., and S.M. wrote the manuscript in consultation with B.F., A.V., T.D., C.A., D.M., W.T., J.M., and S.B,.

## Data availability

The code and data will be made available upon request or acceptance for publishing.

## Supplementary

**Figure 6:**
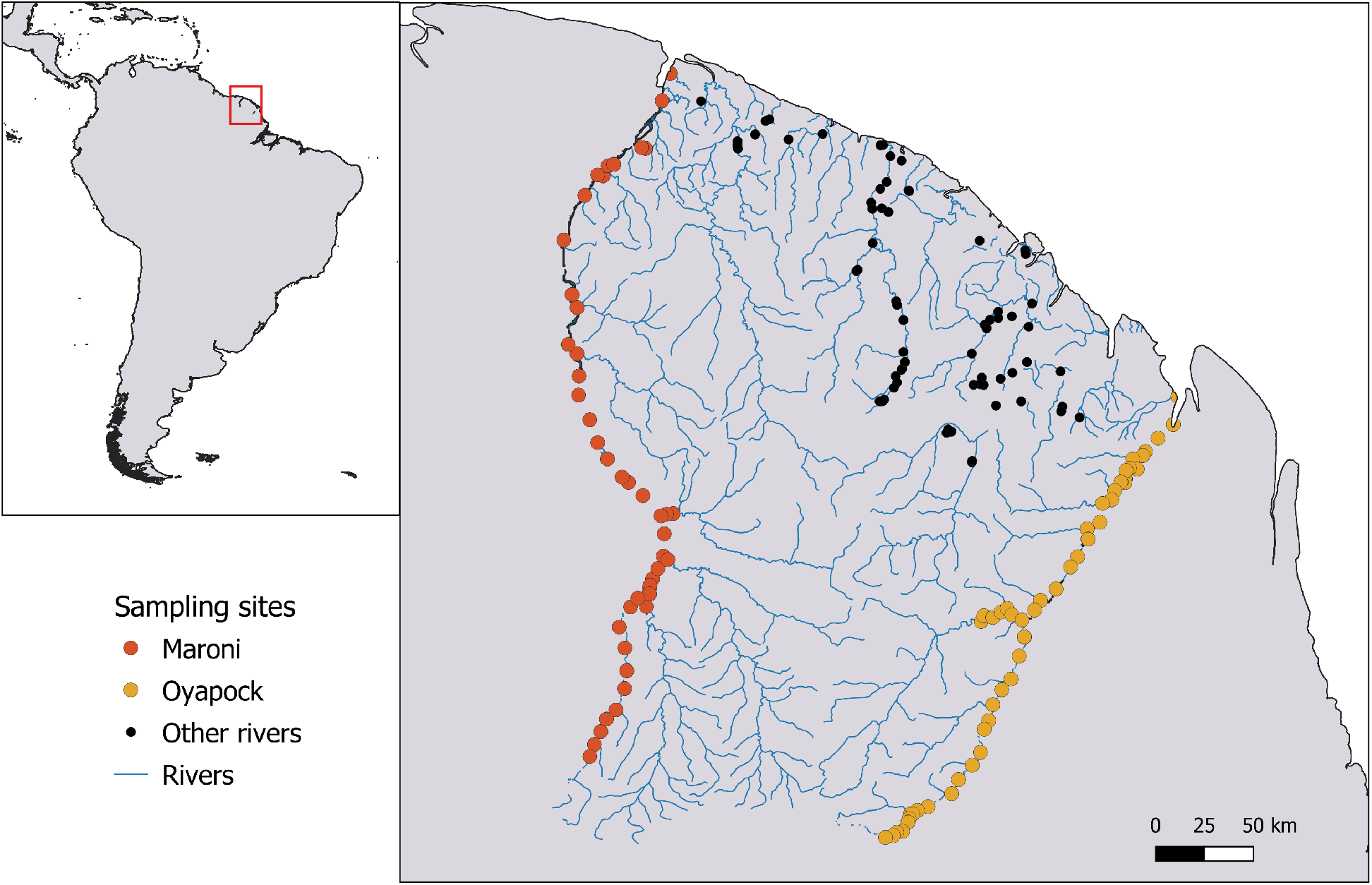
Map of the sampling locations in French Guiana. Sampling sites located on the Maroni river (red), on the Oyapock river (orange), and on other rivers (black).

**Figure 7:**
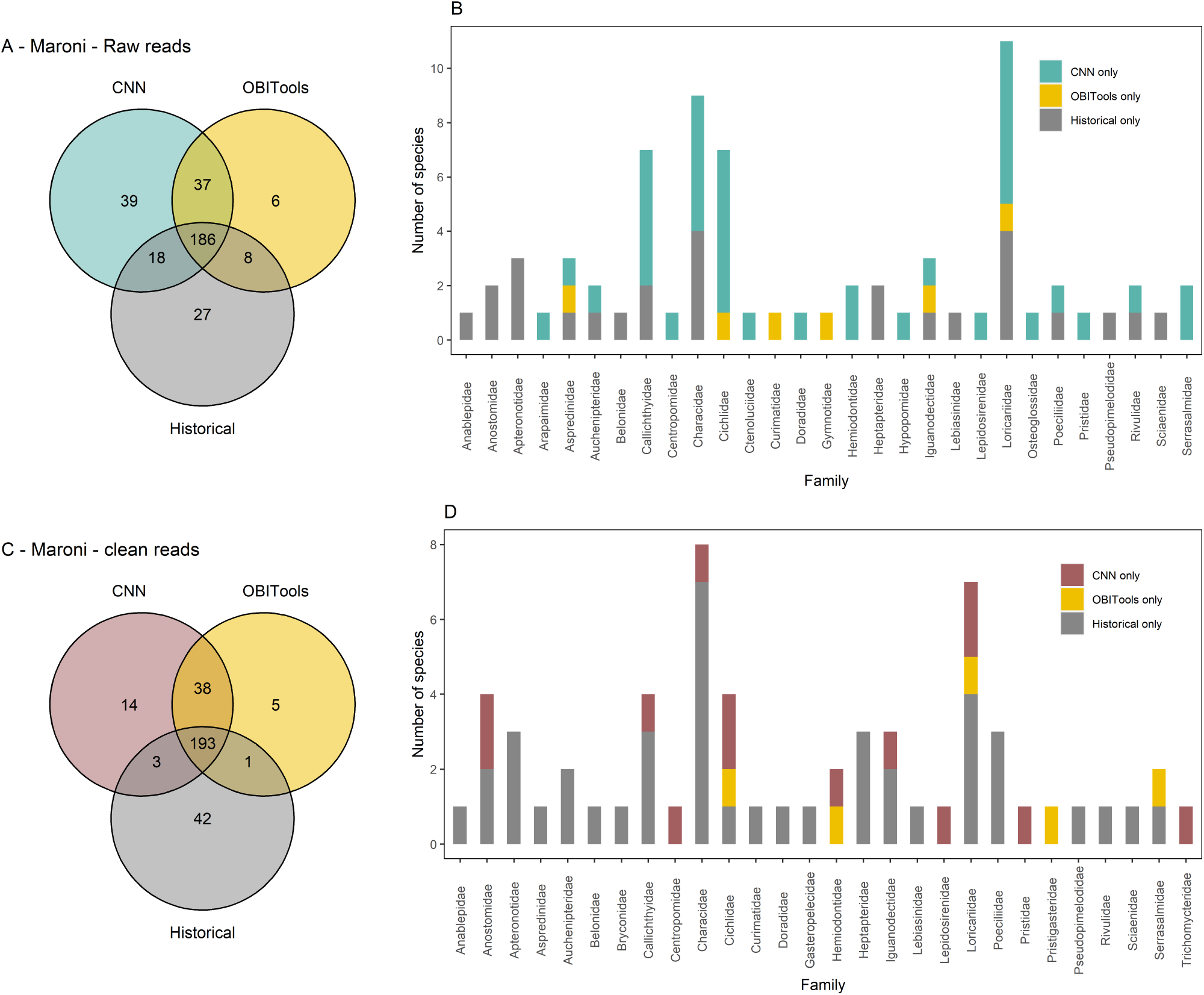
Species detections with the CNN, OBITools and in historical records, in the Maroni river. A, overlap of species detections between the CNN applied on raw reads (blue), OBITools (yellow) and historical records (grey). B, number of species per family, detected with only one method (CNN on raw reads, OBITools or historical records). C, overlap of species detections between the CNN applied on clean reads (red), OBITools (yellow) and historical records (grey). D, number of species per family, detected with only one method (CNN on clean reads, OBITools or historical records).

**Figure 8:**
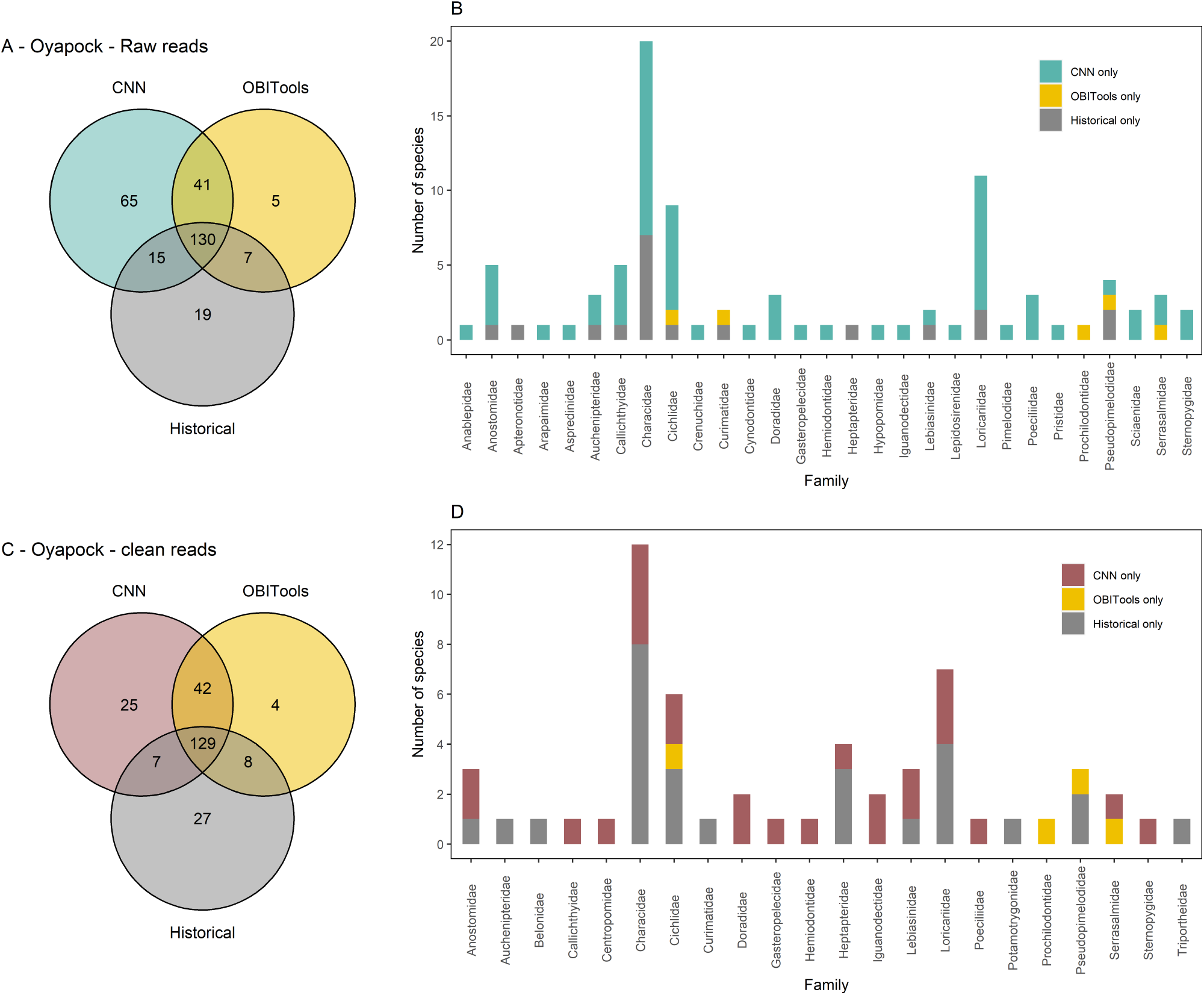
Species detections with the CNN, OBITools and in historical records, in the Oyapock river. A, overlap of species detections between the CNN applied on raw reads (blue), OBITools (yellow) and historical records (grey). B, number of species per family, detected with only one method (CNN on raw reads, OBITools or historical records). C, overlap of species detections between the CNN applied on clean reads (red), OBITools (yellow) and historical records (grey). D, number of species per family, detected with only one method (CNN on clean reads, OBITools or historical records).

**Figure 9:**
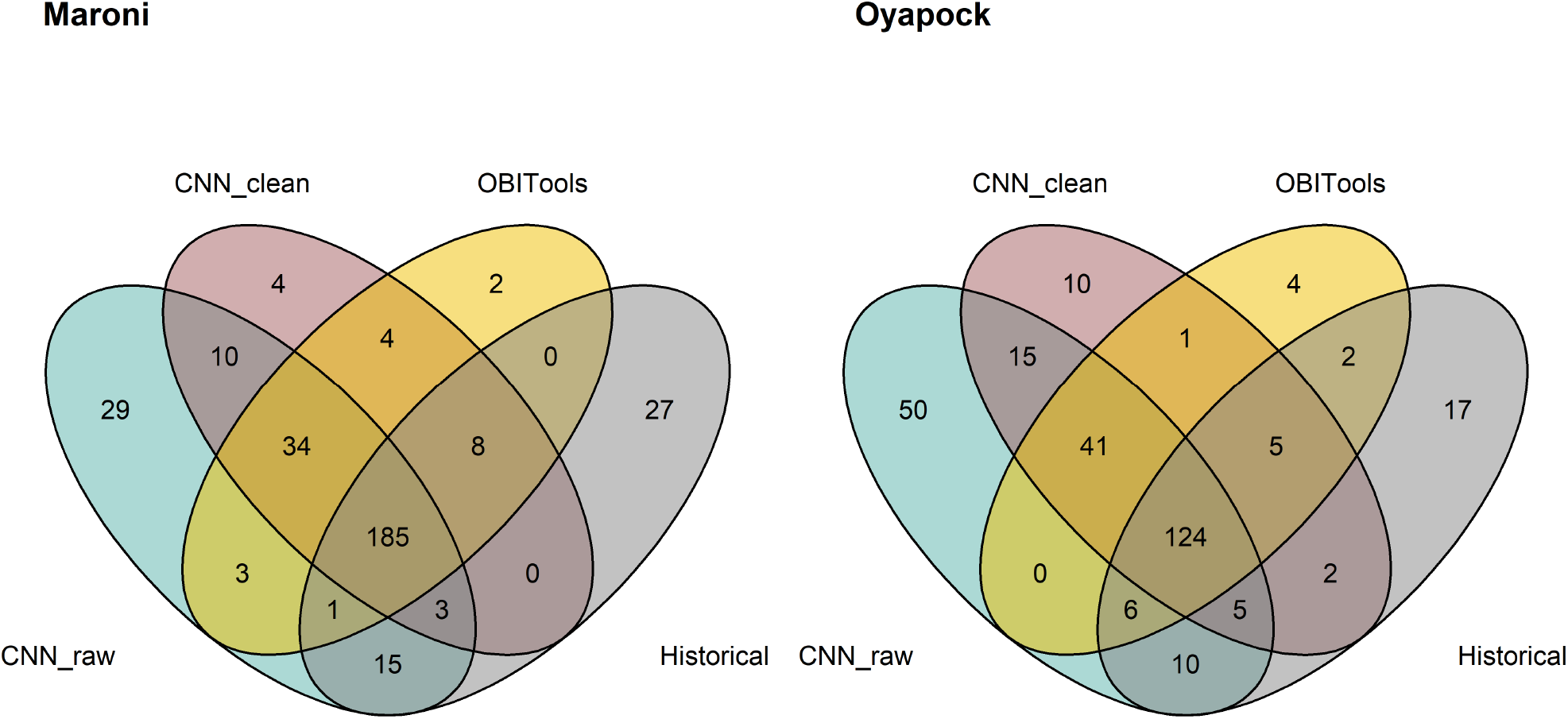
Species detections with the CNN, OBITools and in historical records. Overlap of species detections between the CNN applied on raw reads (blue), the CNN applied on clean reads (red), OBITools (yellow) and historical records (grey). Left for the Maroni river. Right for the Oyapock river.

## References

Abadi, M., Agarwal, A., Barham, P., Brevdo, E., Chen, Z., Citro, C., Corrado, G., … & Zheng, X. (2015). TensorFlow: Large-scale machine learning on heterogeneous systems. Software available from tensorflow.org.

Alberdi, A., Aizpurua, O., Gilbert, M. T. P. & Bohmann K. (2018) Scrutinizing key steps for reliable metabarcoding of environmental samples. Methods Ecology and Evolution. 9, 134–147.

Albert, J. S., & Reis, R. E. (2011). One. Introduction To Neotropical Freshwaters. In Historical biogeography of Neotropical freshwater fishes (pp. 3–20). University of California Press.

Berry, O., Jarman, S., Bissett, A., Hope, M., Paeper, C., Bessey, C., … & Bunce, M. (2020) Making environmental DNA (eDNA) biodiversity records globally accessible. Environmental DNA.

Bohmann, K., Evans A., Gilbert M. T. P., Carvalho, G. R., Creer, S., Knapp, M., Yu Douglas, W., & De Bruyn, M. (2014). Environmental DNA for wildlife biology and biodiversity monitoring. Trends in ecology & evolution 29, no. 6: 358–367.

Bolyen, E., Rideout, J. R., Dillon, M. R., Bokulich, N. A., Abnet, C., Ghalith, G. A. Al, … & Naimey, A. T. (2019). QIIME 2: Reproducible, interactive, scalable, and extensible microbiome data science. Nature Biotechnology, 32, 852–857.

Bonder, M. J., Abeln, S., Zaura, E., & Brandt, B. W. (2012). Comparing clustering and pre-processing in taxonomy analysis. Bioinformatics, 28(22), 2891–2897.

Boussarie, G., Bakker, J., Wangensteen O. S., Mariani, S., Bonnin, L., Juhel, J. B., Kiszka, J. J., Kulbicki, M., Manel, S., Robbins, W. D., Vigliola, L., & Mouillot, D. (2018). Environmental DNA illuminates the dark diversity of sharks. Science Advances 4.

Boyer, F., Mercier, C., Bonin, A., Le Bras, Y., Taberlet, P., & Coissac, E. (2016). obitools: A unix-inspired software package for DNA metabarcoding. Molecular ecology resources, 16(1), 176–182.

Brandt, M.I., Trouche, B., Quintric, L., Günther, B., Wincker, P., Poulain, J., & Arnaud-Haond, S. (2021). Bioinformatic pipelines combining denoising and clustering tools allow for more comprehensive prokaryotic and eukaryotic metabarcoding. Molecular Ecology Resources. Accepted.

Brosse S., Melki F. & Vigouroux R. (2019). Fishes from the Mitaraka mountains (French Guiana). Zoosystema, 41, 131–151.

Brown, E. A., Chain, F. J., Crease, T. J., MacIsaac, H. J., & Cristescu, M. E. (2015). Divergence thresholds and divergent biodiversity estimates: can metabarcoding reliably describe zooplankton communities? Ecology and Evolution, 5(11), 2234–2251.

Busia, K., George, D. E., Fannjiang, C., Alexander, D.H., Dorfman, E., Poplin, R., Chang, P., & DePris, M. (2020). A deep learning approach to pattern recognition for short DNA sequences. BioRxiv.

Bylemans, J., Gleeson, D. M., Hardy, C. M., & Furlan, E. (2018). Toward an ecoregion scale evaluation of eDNA metabarcoding primers: A case study for the freshwater fish biodiversity of the Murray–Darling Basin (Australia). Ecology and evolution, 8(17), 8697–8712.

Calderón-Sanou, I., Münkemüller, T., Boyer, F., Zinger, L., & Thuiller, W. (2020). From environmental DNA sequences to ecological conclusions: How strong is the influence of methodological choices?. Journal of Biogeography, 47(1), 193–206.

Callahan, B. J., McMurdie, P. J., Rosen, M. J., Han, A. W., Johnson, A. J. A., & Holmes, S. P. (2016). DADA2: High-resolution sample inference from Illumina amplicon data. Nature Methods, 13(7), 581–583.

Cantera, I., Decotte, J. B., Dejean, T., Murienne, J., Vigouroux, R., Valentini, A., & Brosse, S. (2020). Characterizing the spatial signal of environmental DNA in river systems using a community ecology approach. bioRxiv.

Cantera, I., Cilleros, K., Valentini, A., Cerdan, A., Dejean, T., Iribar, A., … & Brosse, S. (2019). Optimizing environmental DNA sampling effort for fish inventories in tropical streams and rivers. Scientific Reports, 9(1), 1–1.

Cilleros, K., Valentini, A., Allard, L., Dejean, T., Etienne, R., Grenouillet, G., … & Brosse, S. (2019). Unlocking biodiversity and conservation studies in high-diversity environments using environmental DNA (eDNA): A test with Guianese freshwater fishes. Molecular Ecology Resources, 19(1), 27–46.

Cordier, T., Lanzén, A., Apothéloz-Perret-Gentil, L., Stoeck, T., & Pawlowski, J. (2019). Embracing environmental genomics and machine learning for routine biomonitoring. Trends in microbiology, 27(5), 387–397.

Cordier, T., Alonso-Sáez, L., Apothéloz-Perret-Gentil, L., Aylagas, E., Bohan, D. A., Bouchez, A., & Keeley, N. (2020). Ecosystems monitoring powered by environmental genomics: a review of current strategies with an implementation roadmap. Molecular Ecology.

Coutant, O., Cantera, I., Cilleros, K., Dejean, T., Valentini, A., Murienne, J., & Brosse, S. (2020). Detecting fish assemblages with environmental DNA: Does protocol matter? Testing eDNA metabarcoding method robustness. Environmental DNA.

Deiner, K., Bik, H. M., Mächler, E., Seymour, M., Lacoursière-Roussel, A., Altermatt, F., & Pfrender, M. E. (2017). Environmental DNA metabarcoding: Transforming how we survey animal and plant communities. Molecular ecology, 26(21), 5872–5895.

Deneu, B., Servajean, M., Bonnet, P., Botella, C., Munoz, F., & Joly, A. (2021). Convolutional neural networks improve species distribution modelling by capturing the spatial structure of the environment. PLoS Computational Biology in press.

DiBattista, J. D., Reimer, J. D., Stat, M., Masucci, G. D., Biondi, P., De Brauwer, M., … & Bunce, M. (2020). Environmental DNA can act as a biodiversity barometer of anthropogenic pressures in coastal ecosystems. Scientific reports, 10(1), 1–15.

Dornelas, M., Madin, E. M., Bunce, M., DiBattista, J. D., Johnson, M., Madin, J. S., … & Williams, S. B. (2019). Towards a macroscope: Leveraging technology to transform the breadth, scale and resolution of macroecological data. Global Ecology and Biogeography.

Dufresne, Y., Lejzerowicz, F., Perret-Gentil, L. A., Pawlowski, J., & Cordier, T. (2019). SLIM: a flexible web application for the reproducible processing of environmental DNA metabarcoding data. BMC bioinformatics, 20(1), 1–6.

Ficetola, G. F., Miaud, C., Pompanon, F., & Taberlet, P. (2008). Species detection using environmental DNA from water samples. Biology letters, 4(4), 423–425.

Ficetola, G. F., Taberlet, P., & Coissac, E. (2016). How to limit false positives in environmental DNA and metabarcoding? Molecular ecology resources, 16(3), 604–607

Ficetola, G. F., Pansu, J., Bonin, A., Coissac, E., Giguet-Covex, C., De Barba, M., … & Taberlet, P. (2015). Replication levels, false presences and the estimation of the presence/absence from eDNA metabarcoding data. Molecular ecology resources, 15(3), 543–556.

Flynn, J. M., Brown, E. A., Chain, F. J., MacIsaac, H. J., & Cristescu, M. E. (2015). Toward accurate molecular identification of species in complex environmental samples: testing the performance of sequence filtering and clustering methods. Ecology and evolution, 5(11), 2252–2266.

Gold, Z., Wall, A.R., Curd, E.E., Kelly, R.P., Pentche N.D., Ripma, L., Barber, P.H. & Wetzer, R.(2020). eDNA metabarcoding bioassessment of endangered fairy shrimp (Branchinecta spp.). Conservation Genetics Resources, 12, 685–690.

Grünig, M., Razavi, E., Calanca, P., Mazzi, D., Wegner, J. D., & Pellissier, L. (2021). Applying deep neural networks to predict incidence and phenology of plant pests and diseases. Ecosphere. accepted.

Helaly, M. A., Rady, S., & Aref, M. M. (2019). Convolutional Neural Networks for Biological Sequence Taxonomic Classification: A Comparative Study. In International Conference on Advanced Intelligent Systems and Informatics (pp. 523—533). Springer, Cham.

Holman, L.E., de Bruyn, M., Creer, S., Carvalho, G., Robidart, J., & Rius, M. (2021). Animals, protists and bacteria share marine biogeographic patterns. Nature Ecology & Evolution.

Iknayan, K. J., Tingley, M. W., Furnas, B. J., & Beissinger, S. R. (2014). Detecting diversity: emerging methods to estimate species diversity. Trends in ecology & evolution, 29(2), 97–106.

Jarman, S. N., Berry, O., & Bunce, M. (2018). The value of environmental DNA biobanking for long-term biomonitoring. Nature ecology & evolution, 2(8), 1192–1193.

Juhel, J. B., Utama, R. S., Marques, V., Vimono, I. B., Sugeha, H. Y., Kadarusman, … & Hocdé, R. (2020). Accumulation curves of environmental DNA sequences predict coastal fish diversity in the coral triangle. Proceedings of the Royal Society B, 287(1930), 20200248.

Kopp, W., Monti, R., Tamburrini, A., Ohler, U., & Akalin, A. (2020). Deep learning for genomics using Janggu. Nature communications, 11(1), 1–7.

Le Bail, P. Y., Covain, R., Jégu, M., Fisch-Muller, S., Vigouroux, R., & Keith, P. (2012). Updated checklist of the freshwater and estuarine fishes of French Guiana. Cybium, 36(1), 293–319.

Li, W., Hou, X., Xu, C., Qin, M., Wang, S., Wei, L., Wang, Y., Liu, X. & Li, Y.(2021). Validating eDNA Measurements of the Richness and Abundance of Anurans at a Large Scale. Journal of Animal Ecology. In press.

Makiola, A., Compson, Z. G., Baird, D. J., Barnes, M. A., Boerlijst, S. P., Bouchez, A., & Creer, S. (2020). Key questions for next-generation biomonitoring. Frontiers in Environmental Science, 7.

Marques, V., Guérin, P. É., Rocle, M., Valentini, A. Gold, Z., Wall, A.R., Curd, E.E., Kelly, R.P., Pentcheff N.D., Ripma, L., Barber, P.H. & Wetzer, R. (2020). eDNA metabarcoding bioassessment of endangered fairy shrimp (Branchinecta spp.). Conservation Genetics Resources, 12, 685–690.

Marques, V., Milhau, T., Albouy, C., Dejean, T., Manel, S., Mouillot, D., & Juhel, J. B. (2020b). GAPeDNA: Assessing and mapping global species gaps in genetic databases for eDNA metabarcoding. Diversity and Distributions.

Mathon, L., Valentini, A., Guérin, P. E., Normandeau, E., Noel, C., Lionnet, C.,… & Manel, S. (2021). Benchmarking bioinformatic tools for fast and accurate eDNA metabarcoding species identification. Molecular Ecology Ressources. Accepted

McGee, K. M., Robinson, C., & Hajibabaei, M. (2019). Gaps in DNA-based biomonitoring across the globe. Frontiers in Ecology and Evolution, 7, 337.

Murienne, J., Cantera, I., Cerdan, A., Cilleros, K., Decotte, J. B., Dejean, T., … & Brosse, S. (2019). Aquatic eDNA for monitoring French Guiana biodiversity. Biodiversity data journal, 7.

Nugent, C. M., & Adamowicz, S. J. (2020). Alignment-free classification of COI DNA barcode data with the Python package Alfie. Metabarcoding and Metagenomics, 4, e55815.

Pagni, M., Niculita-Hirzel, H., Pellissier, L., Dubuis, A., Xenarios, I., Guisan, A., … & Guex, N. (2013). Density-based hierarchical clustering of pyro-sequences on a large scale—the case of fungal ITS1. Bioinformatics, 29(10), 1268–1274.

Papa, Y., Le Bail, P. Y., & Covain, R. (2020). Genetic landscape clustering of a large DNA barcoding dataset reveals shared patterns of genetic divergence among freshwater fishes of the Maroni Basin. Authorea Preprints.

Piro, V. C., Dadi, T. H., Seiler, E., Reinert, K., & Renard, B. Y. (2020). ganon: precise metagenomics classification against large and up-to-date sets of reference sequences. Bioinformatics, 36(Supplement1), i12–i20.

Polanco Fernández, A., Marques, V., Fopp, F., Juhel, J. B., Borrero-Pérez, G. H., Cheutin, M. C., Eme, D.,…& Pellissier, L. (2021). Comparing environmental DNA metabarcoding and underwater visual census to monitor tropical reef fishes. Environmental DNA, 3, 142–156.

Polanco Fernández, A., Martinezguerra, M.M., Marques, V., Francisco Villa-Navarro, Borrero-Pérez, G. H., Cheutin, M. C., Dejean, T., Hocdé R., … & Pellissier, L. (2021). Recovering aquatic and terrestrial biodiversity in a tropical estuary using environmental DNA. Biotropica, accepted

Rognes, T., Flouri, T., Nichols, B., Quince, C., & Mahé, F. (2016). VSEARCH: a versatile open source tool for metagenomics. PeerJ, 4, 1–22.

Rojahn, J., Gleeson, D.M., Furlan, E., Haeusler, T.,& Bylemans, J. (2021). Improving the detection of rare native fish species in environmental DNA metabarcoding surveys. Aquatic Conservation: Marine and Freshwater Ecosystems, 31(4), 990–997.

Ruppert, K. M., Kline, R. J., & Rahman, M. S. (2019). Past, present, and future perspectives of environmental DNA (eDNA) metabarcoding: A systematic review in methods, monitoring, and applications of global eDNA. Global Ecology and Conservation, 17, e00547.

Sato, Y., Miya, M., Fukunaga, T., Sado, T., & Iwasaki, W. (2018). MitoFish and MiFish pipeline: a mitochondrial genome database of fish with an analysis pipeline for environmental DNA metabarcoding. Molecular biology and evolution 35.6: 1553–1555.

Schirmer, M., Ijaz, U. Z., D’Amore, R., Hall, N., Sloan, W. T., & Quince, C. (2015). Insight into biases and sequencing errors for amplicon sequencing with the Illumina MiSeq platform. Nucleic Acids Research, 43(6).

Schnell, I. B., Bohmann, K., & Gilbert, M. T. P. (2015). Tag jumps illumi- nated – reducing sequence-to-sample misidentifications in me- tabarcoding studies. Molecular Ecology Resources, 15(6), 1289–1303.

Sepulveda, A. J., Nelson, N. M., Jerde, C. L., & Luikart, G. (2020). Are Environmental DNA Methods Ready for Aquatic Invasive Species Management? Trends in Ecology & Evolution 35, 668–678.

Shokralla, S., Spall, J. L., Gibson, J. F., & Hajibabaei, M. (2012). Next-generation sequencing technologies for environmental DNA research. Molecular ecology, 21(8), 1794–1805.

Singer, G. A. C., Fahner, N. A., Barnes, J. G., McCarthy, A., & Hajibabaei, M. (2019). Comprehensive biodiversity analysis via ultra-deep patterned flow cell technology: a case study of eDNA metabarcoding seawater. Scientific reports, 9(1), 1–12.

Su, G., Logez, M., Xu, J., Tao, S., Villéger, S., & Brosse, S. (2021). Human impacts on global freshwater fish biodiversity. Science, 371(6531), 835.

Taberlet, P., Bonin, A., Coissac, E., & Zinger, L. (2018). Environmental DNA: For biodiversity research and monitoring. Oxford University Press.

Taberlet, P., Coissac, E., Pompanon, F., Brochmann, C., & Willerslev, E. (2012). Towards next-generation biodiversity assessment using DNA metabarcoding. Molecular ecology, 21(8), 2045–2050.

Thomsen, P. F., & Willerslev, E. (2015). Environmental DNA–An emerging tool in conservation for monitoring past and present biodiversity. Biological conservation, 183, 4–18.

Thuiller, W., Lafourcade, B., Engler, R., & Araújo, M. B. (2009). BIOMOD–a platform for ensemble forecasting of species distributions. Ecography, 32(3), 369–373.

Valentini, A., Taberlet, P., Miaud, C., Civade, R., Herder, J., Thomsen, P. F., … & Dejean, T. (2016). Next-generation monitoring of aquatic biodiversity using environmental DNA metabarcoding. Molecular ecology, 25(4), 929–942.

West, K., Travers, M. J., Stat, M., Harvey, E. S., Richards, Z. T., DiBattista, J. D., … & Bunce, M. (2021). Large-scale eDNA metabarcoding survey reveals marine biogeographic break and transitions over tropical north-western Australia. Diversity and Distributions.

